# Evolutionarily diverse LIM domain-containing proteins bind stressed actin filaments through a conserved mechanism

**DOI:** 10.1101/2020.03.06.980649

**Authors:** Jonathan D. Winkelman, Caitlin A. Anderson, Cristian Suarez, David R. Kovar, Margaret L. Gardel

## Abstract

The actin cytoskeleton assembles into diverse load-bearing networks including stress fibers, muscle sarcomeres, and the cytokinetic ring to both generate and sense mechanical forces. The LIM (Lin11, Isl-1 & Mec-3) domain family is functionally diverse, but most members can associate with the actin cytoskeleton with apparent force-sensitivity. Zyxin rapidly localizes via its LIM domains to failing stress fibers in cells, known as strain sites, to initiate stress fiber repair and maintain mechanical homeostasis. The mechanism by which these LIM domains associate with stress fiber strain sites is not known. Additionally, it is unknown how widespread strain sensing is within the LIM protein family. We observe that many, but not all, LIM domains from functionally diverse proteins localize to spontaneous or induced stress fiber strain sites in mammalian cells. Additionally, the LIM domain region from the fission yeast protein paxillin like 1 (Pxl1) also localizes to stress fiber strain sites in mammalian cells, suggesting that the strain sensing mechanism is ancient and highly conserved. Sequence analysis and mutagenesis of the LIM domain region of zyxin indicate a requirement of tandem LIM domains, which contribute additively, for sensing stress fiber strain sites. *In vitro*, purified LIM domains from mammalian zyxin and fission yeast Pxl1 bind to mechanically stressed F-actin networks but do not associate with relaxed actin filaments. We propose that tandem LIM domains recognize an F-actin conformation that is rare in the relaxed state but is enriched in the presence of mechanical stress.

## INTRODUCTION

Cells are subject to a wide range of omnipresent mechanical stimuli, which play essential physiological roles. Epithelial tissue stretch modulates cell proliferation^1,2^, blood pressure regulates the contractility of endothelial cells within blood vessels^3,4^, and muscle contraction shapes connective tissue remodeling^5^. Such mechanotransduction pathways allow for the integration of mechanical cues with the biochemical and genetic circuitry of the cell. While much progress has been made to elucidate the importance of mechanical stimuli in cell physiology, the underlying force-sensing mechanisms and organizational logic of many mechanotransduction pathways are unknown.

To respond to mechanical cues and dynamically modulate cell mechanics, the actin cytoskeleton exploits force-sensitive biochemistry to construct actin filament (F-actin)-based network assemblies. Focal adhesions, the adhesive organelles between cells and their external matrix, can change in composition and size under varied mechanical load^6^. At the molecular scale, these focal adhesion changes arise primarily from force-dependent modulation of constituent proteins^6–8^. The force-dependent association of the focal adhesion proteins, vinculin and talin, to actin filaments is sensitive to filament polarity^9,10^, providing a mechanism to guide local cytoskeletal architecture under load. Protrusive forces at the leading edge of migrating cells are generated by actin polymerization into short-branched F-actin networks^11^. Compressive stress increases the actin filament density, which, in turn, alters its force generation potential^12^. At the molecular scale, this mechanical adaptation of lamellipodial networks may arise from increased branch formation efficiency by the actin nucleator Arp2/3 complex on extensionally strained sides of bent actin filaments^13^. Furthermore, the affinity of the severing protein cofilin may be lower for taut actin filaments^14^, which reduces filament disassembly if they are under tension^15,16^. While the mechanism by which these proteins sense mechanical deformations of actin filaments is still unknown, the structural polymorphism in actin filaments strongly suggests the likelihood of force-induced conformations^17^. Thus, force-modulated affinity of binding partners with actin filaments is likely widespread within the actin cytoskeleton as a means to regulate cell mechanotransduction pathways.

Within adherent cells, F-actin bundles known as stress fibers (SFs) generate contractile force across the cell and, via focal adhesions, are coupled to the extracellular matrix. SFs dynamically rearrange over long (hour) time scales in response to forces applied to the extracellular matrix^18^, and this remodeling process requires zyxin and paxillin^18–22^. At short times, mechanical failure of the SF can occur either spontaneously or in response to applied force^19,21^. At such damage sites, the SF is locally weaker, leading to a localized retraction to create a stress fiber strain site (SFSS)^21^ (Fig. 1a). Rather than irreversible failure, a repair process is initiated at SFSS that is initiated by the rapid accumulation of zyxin and followed by the recruitment of binding partners a-actinin and VASP to promote actin assembly and cross-linking to repair the SF and maintain mechanical homeostasis^21^ (Fig. 1a). The recruitment of a-actinin and VASP require known interactions at the zyxin N-terminus^22^. However, the recruitment of zyxin to SFSS occurs through a region near the C-terminus that contains three sequential <u>L</u>in11, <u>I</u>sl-1 & <u>M</u>ec-3 (LIM) domains separated by two short 7-8 residue-length unstructured linkers (Fig. 1b,c)^21^. While this LIM domain-containing region (LCR) is necessary and sufficient for localization to SFSS, the underlying mechanism is not known. It has been speculated that the signal within SFSS that the LCR senses may arise from new F-actin barbed ends, conformational changes to an actin binding protein, or post-translational modifications of actin or zyxin’s binding partners^22^. Moreover, the focal adhesion protein paxillin also localizes to SFSS through its LCR^19^, suggesting that this apparent mechanosensing process may be more generally conserved.

**Figure 1.**
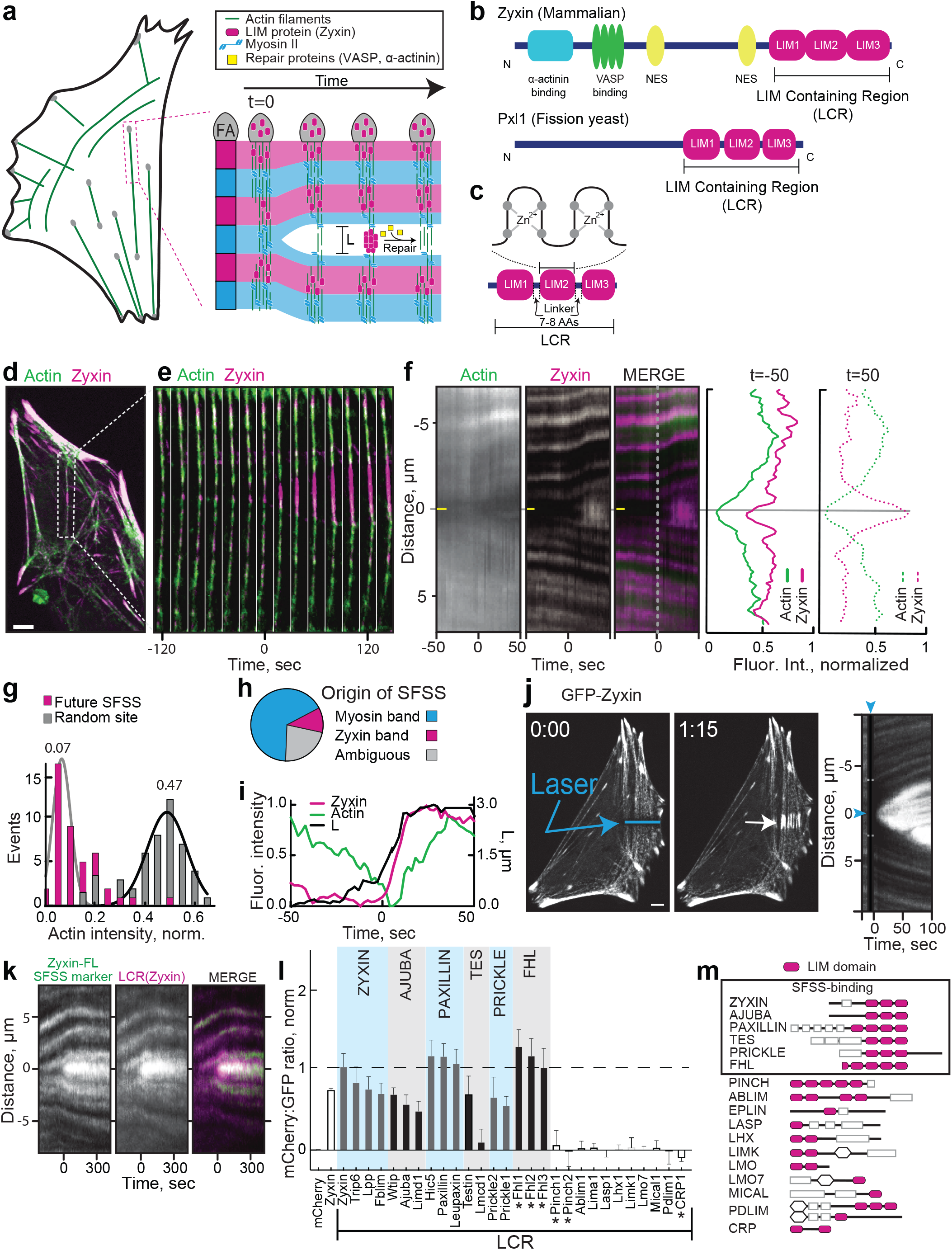
Diverse LIM domains localize to SFSS. (a) Cartoon of a fibroblast cell with actin stress fibers. Schematic of the development and repair of a stress fiber strain site (SFSS). FA=Focal adhesion. L is distance across the SFSS. (b) Domain organization of the LIM domain (pink ovals)-containing proteins mammalian zyxin and fission yeast Pxl1. Zyxin also contains binding sites for α-actinin (F-actin crosslinker) and VASP (F-actin elongation factor), and two nuclear export sequences (NES). LIM containing regions (LCR) are indicated. (c) Each LIM domain contains two zinc finger binding domains with conserved residues (cysteine/histidine, grey circles) to chelate the zinc, but the remaining sequence varies between different LIM domains. The linker length between adjacent LIM domains is 7-8 amino acids. (d-i) Analysis of SFSS in mouse embryo fibroblasts (MEF) with stably integrated GFP-zyxin and transfected mApple-actin. Scale bar=5 μm. (d-e) Fluorescent micrograph (d) and associated montage (e) of a representative stress fiber over time showing accumulation of zyxin on a developing SFSS. (f, left) Kymographs of the same event: actin channel (left), zyxin channel (middle), and a merged image (right). The future SSFS (horizontal yellow line) indicates where background measurements for (g) and a later screen will be taken. Vertical dotted line indicates when strain has begun at t=0. (f, right) Average fluorescence intensity line scans of 74 SFSS, measured 50 seconds before and after initiation. (g) Histogram of actin intensities at a future SFSS (~50 seconds prior) and random sites on the same stress fiber. (h) Pie chart showing the distribution of where SFSS occurred. (i) Kinetics of zyxin accumulation and actin depletion and reassembly (left y-axis), and distance across the SFSS indicated as L in (a) (right y-axis), for a representative SFSS. (j) Left, fluorescent micrographs showing laser induction of a SFSS in a representative MEF cell with stably integrated GFP-zyxin. Blue line shows where light was targeted, and white arrow denotes developing SFSS. Scale bar=5μm. Right, kymograph showing this event over time, with blue arrowheads indicating time and location of laser light. (k) Representative kymograph of the laser-induced SFSS screen, from a cell expressing GFP-zyxin and LCR(zyxin)-mCherry. (l) Screen of 28 LIM domain proteins from Mus musculus. Y-axis is the mCherry:GFP ratio at the strain site, error bars=95% CI. LCR constructs were used for all but those marked with *, for which the whole protein sequence were used. (m) Domain organizations of LIM families in mammals. Box denotes the families that bound to SFSS.

The family of proteins that contain one or more LIM domains is large. In humans, there are ~70 genes containing LIM domains that are divided into 14 families^23^, many of which associate with load bearing elements of the cytoskeleton, such as focal adhesions, cell-cell adhesions, and stress fibers^22^. There are at least 26 LIM domain-containing proteins that localize to focal adhesions, many of which require cell contractility for proper localization^7,8^. While LIM domain-containing proteins are ubiquitous in diverse mechanotransduction pathways, it is unknown whether these share a mechanism by which mechanical stimuli is transduced. Moreover, the mechanism by which the LCR is recruited to SFSS is unknown.

We employed a combination of live cell imaging and *in vitro* reconstitution approaches to explore cytoskeletal mechanosensing by LIM domain proteins. We first established a SFSS-localization screen of diverse LCRs from both mammals and yeast and identified 17 LCRs within the testin, prickle, four and a half LIMs (FHL), paxillin, and zyxin protein families that bound to SFSS. We also determined that paxillin-like 1 (Pxl1) from fission yeast localizes to SFSS in mammalian cells, suggesting that the strain activated target produced in SFSS and recognized by LCRs is well conserved. Sequence and domain analysis of SFSS-binding LCRs shows that tandem LIM domains contribute additively to SFSS association. To identify the strain activated target of LCR, we reconstituted contractile actomyosin networks *in vitro* and observed LCR localization to mechanically stressed actin filaments. From these data, we propose that tandem LIM domains bind a periodic molecular mark that emerges on strained actin filaments.

## RESULTS

### LIM-domain containing regions (LCRs) from diverse mammalian proteins bind to SFSS

SFs are contractile bundles of 10-30 crosslinked actin filaments with alternating bands enriched with either myosin II motor or α-actinin, VASP, and zyxin (Fig. 1a)^22^, a structure similar to that in striated myofibrils^24^. SFSS develop when SFs mechanically fail, resulting in local elongation and thinning that compromise their force transmission^21^ (Fig. 1a, d-f). Zyxin rapidly accumulates at the SFSS and recruits the actin assembly factor VASP and crosslinking protein α-actinin to repair and stabilize the damaged site (Fig. 1a)^21^. Previous work identified the LCR of zyxin to be necessary and sufficient for accumulation on SFSS^19^. Measuring the fluorescence intensity of zyxin and actin along the SF prior to a SFSS reveals locally diminished intensity of both proteins at the future SFSS (Fig. 1d-f). At the initial stages of repair just after *t*=0 s, zyxin rapidly intensifies at the location of minimal actin (Fig. 1d-f). Examining the local actin intensity of a future SFSS, we find that the actin intensity is depleted five-fold as compared to regions of SFs that do not fail (Fig.1f,g and Supplementary Video 1). Moreover, we find that >65% of SFSS occur in a myosin-rich band (Fig. 1h), and myosin II is displaced laterally from SFSS (Supplementary Fig.1a-c). The filament density decreases as the SFSS expands^21^, and zyxin recruitment occurs nearly simultaneously as the length *L* of the strain site increases (Fig. 1i). These data suggest that SFSS occur at SF regions pre-disposed to failure because of lower actin density and depletion of actin assembly and cross-linking factors. We also found that SFSS can be induced by partially damaging the SF with high laser intensity (Fig. 1j and Supplementary Video 2). The LCR of zyxin is recruited to laser-induced SFSS with similar kinetics to that of spontaneous SFSS (Fig. 1k and Supplementary Fig. 1d). Considering force balance along the SF in either of these two scenarios, the reduced number of actin filaments at SFSS suggests filaments and crosslinks present there are under an increased load.

To assess whether localization to SFSS is a feature ubiquitous within the LIM family of proteins, we developed an assay to quantify their recruitment to either endogenous or induced SFSS. We cloned the LCR from one or more genes belonging to each LIM protein class^23^ and generated mCherry-tagged mammalian expression constructs. Each of the 28 mCherry-tagged LCRs was transiently transfected into mouse embryo fibroblast (MEF) cells with GFP-zyxin stably integrated into the genome. Using the GFP-zyxin as a positive marker for SFSS, we then assessed the localization of the mCherry-tagged LCRs at the site (Fig. 1k). As a control, the mCherry-tagged constructs of both full length zyxin and the LCR of zyxin, LCR(zyx), localize very similarly to GFP-zyxin (Fig. 1k,l and Supplementary Video 3). At a SFSS, we generated kymographs (Fig. 1k) and, from these, took linescans across the time axis at the center of the SFSS to generate a kinetic profile of SFSS accumulation for both GFP-zyxin and the transfected LCR-mCherry. From these profiles, the ratio of the LCR-mCherry to GFP-zyxin was determined and normalized (Supplementary Fig. 1e-n) such that a cytoplasmically expressed, mCherry-tagged nuclear export signal (NES) which was added to all LCRs, is zero and mCherry-LCR(zyx) is one (Fig. 1l and Supplementary Fig. 1n). Ratio averages that were significantly above zero were scored as SFSS-binders. We observed SFSS localization of the LCRs from 16 proteins across several LIM protein classes, including zyxin, ajuba, paxillin, testin (Tes), prickle, and four and a half LIMs (FHL) (Fig. 1l,m). SFSS-sensing is isolated to, but ubiquitous within, these classes as the LCRs of all but one protein tested from these classes (Lmcd1) localize to SFSS significantly above background. These results identify novel LIM domain protein-sensitivity to mechanical strain in the actin cytoskeleton and demonstrate its conserved function across diverse LIM domain containing proteins.

### LIM Domains from fission yeast bind to SFSS in mammalian cells via a conserved mechanism

To explore when along evolutionary lines SFSS-binding in LIM domain proteins might have arisen and to determine the level of conservation, we looked for SFSS-binding homologues in more divergent species. The paxillin family first appears in the unikonts (amoebas, yeasts, metazoans) while testin, prickle, FHL, zyxin, and ajuba families arose later in the metazoans^25^. Reflecting a much simpler genome, fission yeast express five LIM domain-containing proteins: Rga1, Rga3, Rga4, Hel2, and Pxl1 (Fig. 2a). The only contractile actin filament network in fission yeast is the cytokinetic ring, where myosin rich nodes condense into an actomyosin bundle that constricts to drive cell division^26^. Time-lapse imaging of fission yeast Pxl1-GFP shows strong co-localization with myosin II at the contractile ring but only after the ring has assembled and begins to constrict (Fig. 2b)^27,28^. Although localization to the constricting contractile ring may suggest mechanosensitive localization, we could not manipulate the mechanics of the ring to directly test this possibility.

**Figure 2.**
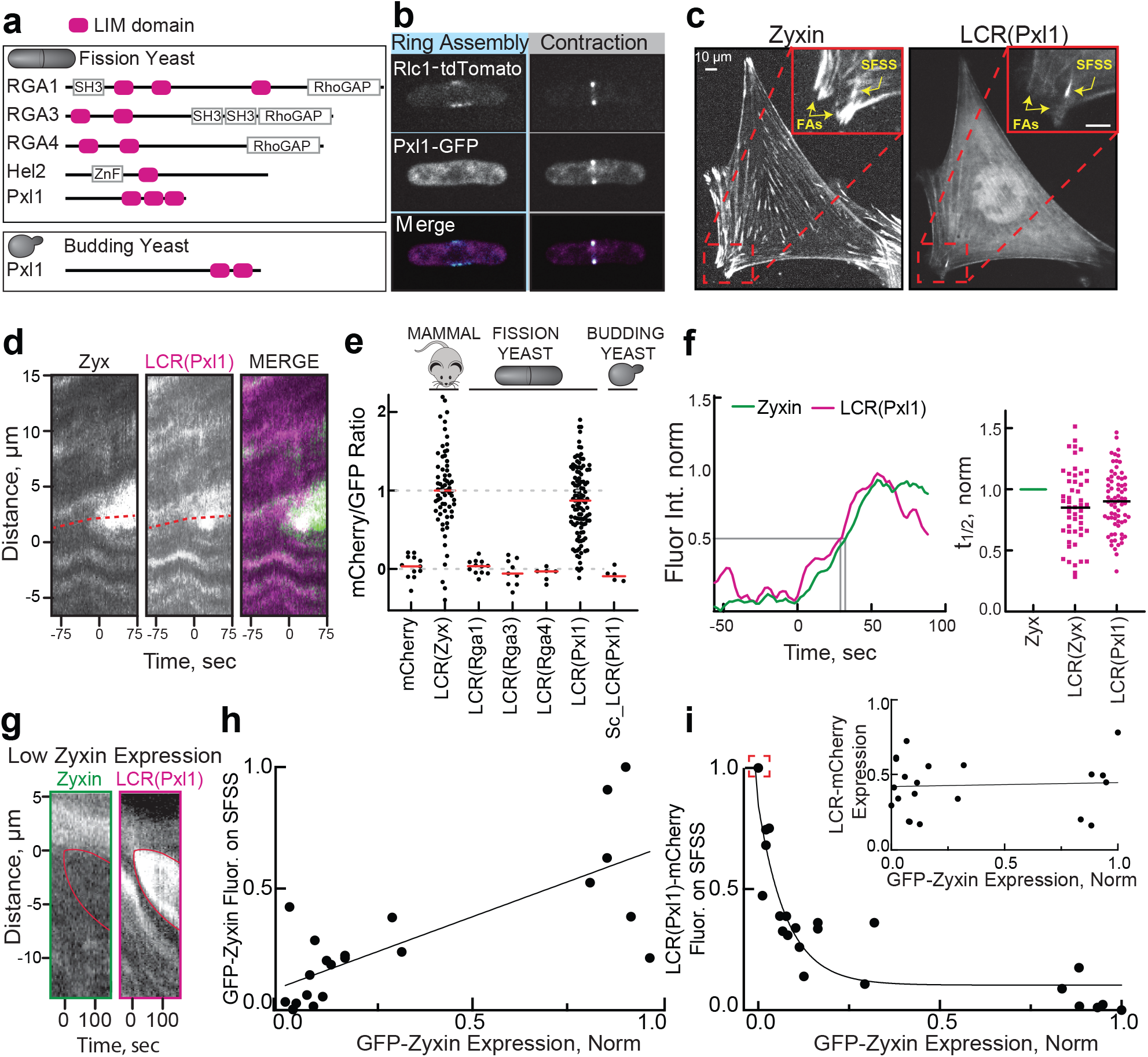
LCR of fission yeast Pxl1 associates to SFSS and displays similar kinetics and competes at SFSS. (a) Domain organization of *Schizosaccharomyces pombe* (fission yeast) LIM domain proteins and budding yeast Paxillin-like 1 (Pxl1). (b) Fluorescent micrographs of a fission yeast cell undergoing cytokinesis that is expressing myosin II regulatory light chain Rlc1-tdTomato and full length Pxl1-GFP. Contractile ring assembly (left panels) and beginning of contraction (right panels) are shown. (c) Mouse embryo fibroblast (MEF) cell expressing GFP-zyxin and the LIM domains of fission yeast Pxl1, LCR(Pxl1)-mCherry. Focal adhesions (FAs) and SFSS (yellow arrows) are labeled in the magnified inset image. (d) Kymograph of SFSS in the MEF cell from (c). (e) Screen for SFSS-association in fission yeast and budding yeast LCRs. (f, left) Plot of the accumulation of GFP-zyxin and fission yeast LCR(Pxl1)-mCherry on SFSS over time. Grey lines indicate t_1/2_ and t_max_ for GFP and mCherry. (f, right) Plot of the normalized t_1/2_ for each SFSS. LCR(Zyx)- or LCR(Pxl1)-mCherry was normalized to GFP-zyxin, which was set to one. LCR(Pxl1)-mCherry t_1/2_ distribution centered around 0.97, st. dev.=0.27, n=120 SFSSs from >10 cells. (g-i) Fluorescence intensity of zyxin and LCR(Pxl1) at SFSS correlates with expression levels. (g) Kymograph showing competition of fission yeast LCR(Pxl1)-mCherry with GFP-zyxin at a MEF cell SFSS. Expression levels were estimated by measuring cytoplasmic intensity. (h) Fluorescence signal of fission yeast LCR(Pxl1)-mCherry or GFP-zyxin at SFSS as a function of estimated expression levels of the same construct. R^2^=0.57, n=21. (i) Fluorescence signal of LCR(Pxl1) at SFSS as a function of estimated expression of GFP-zyxin. R^2^=0.81. (i, inset) Estimated expression of LCR(Pxl1)-mCherry as a function of GFP-zyxin expression. R^2^=0.003.

To determine whether SFSS localization found in mammalian LCRs is preserved in the fission yeast LIM domain-containing proteins, we used the SFSS-localization assay developed for mammalian cells. We transfected fission yeast LCRs tagged with mCherry into MEF cells containing stably integrated GFP-zyxin. The LCR of fission yeast Pxl1, LCR(Pxl1), localizes with the periodic z-bands in SFs but is largely absent from focal adhesions (Fig. 2c). Surprisingly, LCR(Pxl1) exhibits strong SFSS localization, similar to that observed with LCR(zyx) (Fig. 2c-e and Supplementary Video 4). Conversely, the LCR of fission yeast RhoGAPs and budding yeast Pxl1 did not display SFSS localization (Fig. 2e). To determine if the fission yeast LCR(Pxl1) binds to the same target in SFSS as mammalian LCR(zyx), we compared the kinetics of LCR(Pxl1) and zyxin accumulation at SFSS (Fig. 2f) and calculated the time to reach half of the maximum fluorescence intensity (t_½_). The t_½_ of LCR(Pxl1) is nearly identical to LCR(zyx) (Fig. 2f), strongly suggesting that the two highly divergent LCRs use the same mechanism for SFSS association.

As a second test of whether LCR(Pxl1) and LCR(zyx) sense the same binding site to associate with SFSS, we assayed whether they compete for association to the same SFSS. For these experiments, we exploited natural variations in the expression of zyxin-GFP and LCR(Pxl1)- mCherry in our cell populations (Fig. 2g-i). As expected, cells expressing high levels of zyxin showed more zyxin signal at SFSS (Fig. 2h), which is also true for LCR(Pxl1) (Supplementary Fig. 2b). We also verified that expression of zyxin is not correlated with expression of LCR(Pxl1) (Fig. 2i, inset). Importantly, high zyxin expression inversely correlates with reduced SFSS association of LCR(Pxl1) (Fig. 2h,i). For instance, when expressed at low levels, zyxin accumulation at SFSS is nearly completely inhibited by LCR(Pxl1) (Fig. 2g,i), and conversely, very little LCR(Pxl1) accumulates at SFSS in cells expressing high levels of zyxin (Fig. 2i). The competitive relationship between zyxin and LCR(Pxl1) argues that these diverse LCRs (Supplementary Fig. 2c,d) are recruited to SFSS via the same mechanism. Although yeasts may not have canonical SFs, the contractile ring may still exhibit processes similar to SFSS^28,29^ that are recognized by Pxl1. Thus, although fission yeast do not contain SFs, a highly conserved molecular feature that exists in both fission yeast and mammalian SFSS is recognized by LCR(Pxl1). We conclude that the target of SFSS-sensing LCRs existed in the common ancestor of yeast and mammalian cells, and this association of LCRs with the actin cytoskeleton has likely been conserved since at least the divergence of yeasts and mammals.

### Tandem LIM domains contribute additively to SFSS localization

Since LCRs from mammals and fission yeast appear to recognize a common target in SFSS, we speculated that conserved amino acid sequence signatures may be required for this function. However, no obvious signature is revealed by comparing the alignment of LIM domain amino acid sequences of mammalian SFSS-binders and non-binders (Supplementary Fig. 2c). Outside of the highly conserved amino acids that coordinate the zinc ions (Cys, His or Asp), the sequences of individual LIM domains are highly variable. Within the LCR of zyxin, there are three LIM domains (LIM1, LIM2, LIM3) separated by 2 short linkers (Fig. 1c). LIM2 and LIM3 share only 33% and 26% sequence identity to LIM1 (Fig. 3a). Furthermore, LCRs from SFSS-binding and non-binding proteins show similar level of sequence identity (30% and 28%) to LIM1 of zyxin (Fig. 3a). However, we noticed common organizational elements of the LCR in SFSS-binders. First, all SFSS-binders contain three or more LIM domains in tandem organization, while 85% of non-binders have fewer than three LIM domains (Fig. 3b). For all proteins in prickle, testin, FHL, paxillin, and zyxin classes, we plotted the level SFSS binding determined from the screens in (Fig. 1m, 2e) versus the number of LIM domains within the LCR. We found that the degree SFSS binding is weakly correlated (R^2^ = 0.7) with the number of tandem LIM domains (Fig. 3c). Furthermore, linkers from SFSS-binding proteins are 7-8 amino acids long, while linkers in non-binders range widely from 7-200 amino acids (Fig. 3d). These findings suggest a testable hypothesis that tandem LIM domains connected by a short linker is necessary for SFSS binding (Fig. 3e).

**Figure 3.**
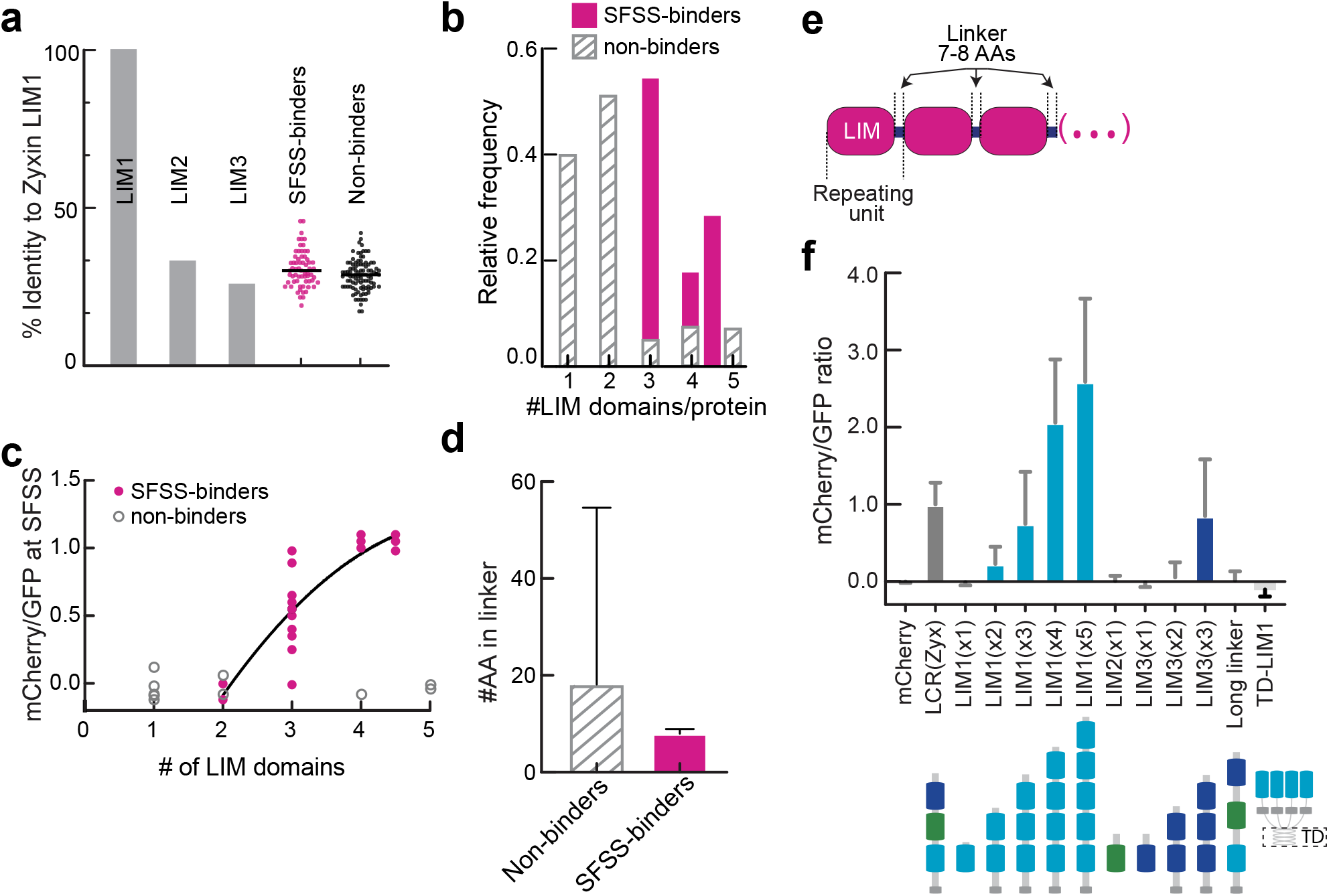
LCRs bind to SFSS through multiple, precisely spaced domains organized in tandem. (a) Amino acid identity to mammalian zyxin’s first LIM domain (LIM1) of zyxin’s other LIM domains (LIM2 and LIM3) and LIM domains from SFSS-binder and non-binder families. (b) Histogram of the number of individual LIM domains in each SFSS-binder and non-binder protein. (c-e) Common organizational elements of LCRs in SFSS-binders. (c) Plot of SFSS-binding vs the number of LIM domains in the LCR construct. Data from LCRs within LIM binding classes was fit with a quadratic function using least squares. R^2^=0.68. (d) Lengths of linkers (in amino acids) connecting LIM domains. Error bars=st.dev. (e) Domain organization of a typical SFSS-binding LCR. There are typically 3 (or more) LIM domains separated by linkers of 7-8 residues. (f) SFSS-association screen of various organizations of zyxin’s LCR. LIM1(X3) indicates the LIM1 is repeated 3 times. TD-LIM1 indicates the LIM1 domain was oligomerized by the addition of a modified GCN4 tetramerization domain (TD).

We then explored how alterations to LCR(zyx) organization impacts its localization to SFSS using the screening approach described previously. Constructs containing any one LIM domain of zyxin (LIM1(x1), LIM2(x1), or LIM3(x1)) do not localize to SFSS (Fig. 3f). However, linkage of multiple LIM1 domains in tandem connected by a linker 8 AA long (See materials and methods) are recruited to SFSS (Fig. 3f). Two tandemly linked LIM domains (Lim1(X2), LIM3(X2) and LIM1LIM2) weakly associate with SFSS (Fig. 3f, Supplementary Fig. 3a,b). Additionally, three tandemly linked LIM domains (LIM1(x3)) or LIM3 (LIM3(x3)), exhibit SFSS localization similarly well as LCR(zyx) (Fig. 3f). When the number of LIM1 repeats is increased to four (LIM1(x4)) or five (LIM1(x5)), the localization to SFSS further increases, exceeding LCR(zyx) by up to two-fold (Fig. 3f). These data suggest that each LIM domain alone may weakly bind the target within SFSS, but multiple interactions (at least for LIM1 and LIM3) contribute to the avidity of target binding. To determine whether the specific organization of LIM oligomerization matters, we clustered four LIM1 in parallel with a synthetic GCN4 tetramerization domain (TD) that drives the formation of a left handed coiled coil^30^, TD-LIM1(x4). However, we did not observe SFSS localization with this construct (Fig. 3f) or from GCN4 dimerization and trimerization domains (Data not shown). Since the length of linkers in SFSS-binding LCRs is highly conserved (Fig. 3d), we tripled the linker lengths in LCR(zyx) from 8 to 24 amino acids (Long Linker), which abrogated SFSS localization (Fig. 3f). Thus, full binding of LCR to the target within SFSS requires at least three tandem LIM domains connected with a short linker. This supports the notion that serial organization of multiple LIM domains is important for SFSS localization.

### In vitro Reconstitution of LIM recruitment to contractile actomyosin bundles

While the above experiments identify organizational features of LCR required for association with SFSS, its binding target remains unclear. These experiments also indicate that LCR binding to SFSS is highly conserved from mammals to fission yeast, suggesting the binding target of LCR may be a core component of the eukaryotic contractile machinery. To identify the target within SFSS that is recognized by LCRs, we chose an *in vitro* reconstitution approach with a well-studied set of purified proteins. Untagged and SNAP-tagged LCR proteins SNAP-LCR(Pxl1) and SNAP-LCR(zyx) (Fig. 4a, Supplementary Fig. 4a) were purified from bacteria, with modified expression conditions to optimize for stability of Zn-coordinated protein, and the SNAP tag was fluorescently labeled with SNAP-surface 549 or 647 for single molecule imaging using total internal reflection fluorescence (TIRF) microscopy. Untagged and SNAP-tagged LCR(zyx) and LCR(Pxl1) elute from a gel filtration column as a stable monomer.

**Figure 4.**
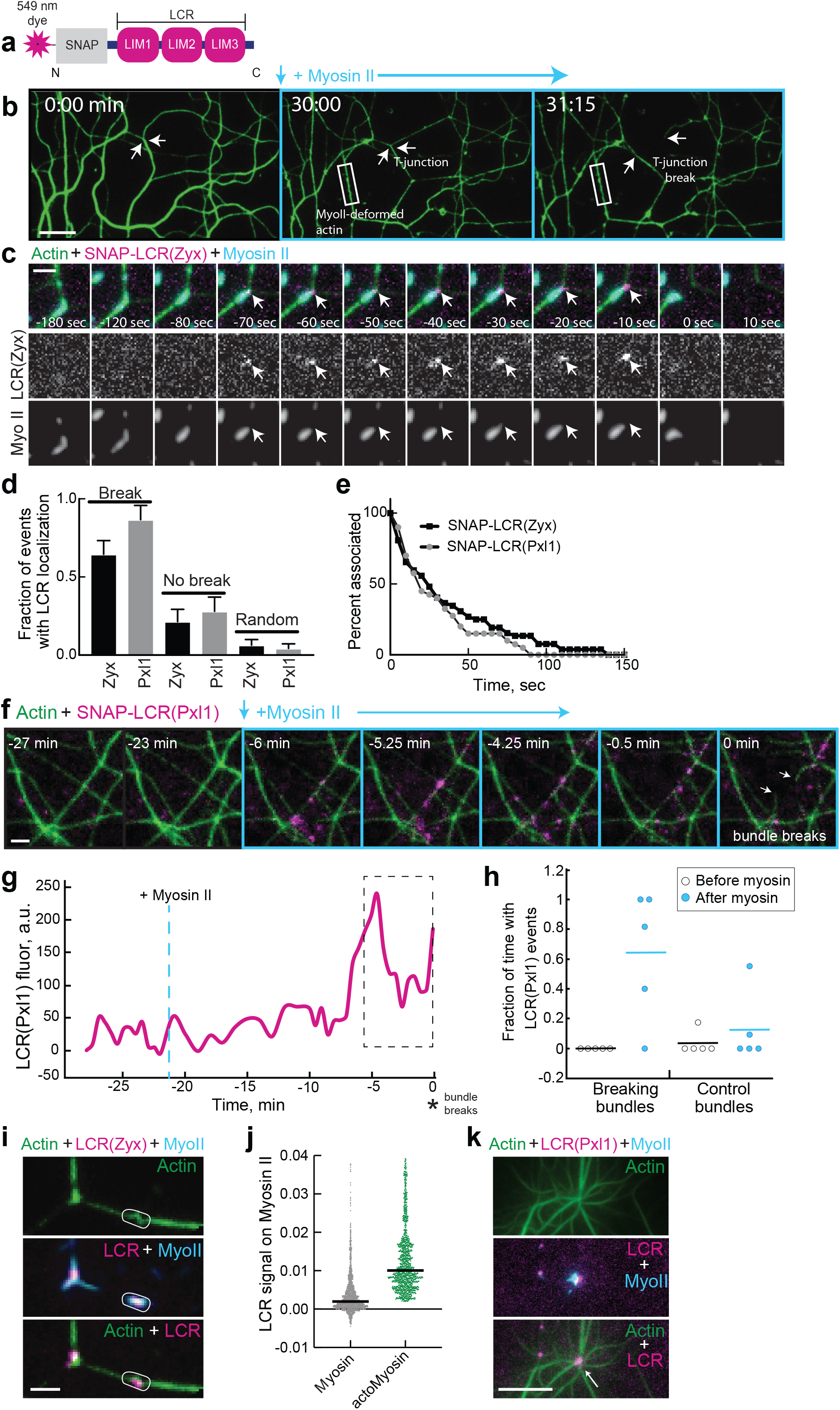
Purified LCR of yeast Pxl1 and mammalian zyxin localized to stressed F-actin networks. (a) Schematic of the SNAP-tagged LCR protein constructs used for in vitro experiments. (b-k) TIRFM visualization of the recruitment of LCR protein constructs to actin filament networks. F-actin networks were preassembled with Mg-ATP-actin (10% Alexa488-labeled), α-actinin and the indicated LCR construct (SNAP-549-tagged,) for 30-45 minutes. Network contraction was subsequently induced by flowing in polymerized myosin II with actin (0.1 μM) and the same initial concentrations of α-actinin and LCR. (b) A representative time-lapse of the in vitro contraction assay. After myosin is added to the preassembled bundled F-actin network, several types of network stresses occur, including T-junctions (white arrow) and myosin-induced F-actin deformations (white box). Scale bar=10 μm. (c-e) LCR localization to T-junctions. (c) Representative time-lapse montage. A preassembled network was formed with 1.5 μM actin, 75 nM α-actinin, and 100 nM SNAP-LCR(zyx), followed by the addition of 100 nM myosin to induce network contraction. White arrows show when LCR(zyx) localizes to the T-junction prior to break. Scale bar =2 μm. (d) Quantification of LCR association at T-junctions. LCR was either present (1) or absent (0) at T-junctions in the frame before breaking, T-junctions that did not break, or at random sites along the bundle. Error bars represent SEM, n=individual T-junctions or random site. (e) Lifetime of LCR on T-junctions that broke, which displayed single molecule fluorescence intensity. (f-h) LCR localization to less common breaks along filament bundles. (f) Representative time-lapse montage of a preassembled network formed with 3.0 μM actin, 150 nM α-actinin, and 200 nM SNAP-LCR(Pxl1), followed by the addition of 75 nM skeletal muscle myosin II to induce network contraction. Scale bar=2 μm. White arrows indicate broken ends of a bundle. (g) LCR(Pxl1) recruitment measured along the F-actin bundle in (f) that breaks at time 0. (h) LCR(Pxl1) fluorescence along bundles was measured similarly as in (g) for 5 bundle breaking events. Since LCR recruitment appears to be dynamic along the bundle, the fraction of frames with LCR localization during the 5 minutes just prior to break was measured. (i-j) LCR localization to myosin along stressed F-actin networks, particularly where myosin deforms F-actin bundles. (i) Representative fluorescent image of LCR(zyx) localizing with myosin II along the network. Ovals indicate myosin on the F-actin network or on the glass. (j) Quantification of LCR localization with myosin or myosin associated with actin (actomyosin). (k) Representative fluorescent images of a large aster that forms from a preassembled network with 3.0 μM actin and 200 nM SNAP-LCR(Pxl1) in the absence of α-actinin, followed by the addition of 90 nM myosin II (Alexa 647 labeled). Scale bar=10 μm. LCR(Pxl1) localizes to the center of these asters with the myosin (white arrow).

A leading hypothesis is that LCR(zyx) binds to actin filament barbed ends produced from filament breakage in SFSS^21^. We first tested whether LCR(zyx) or LCR(Pxl1) bind to actin filament sides or barbed ends in standard bulk assays (Supplementary Fig. 4a,b). Actin filament sedimentation assays revealed a barely detectable increase of LCR in the pellet over a range of increasing actin filament concentrations, indicating an extremely weak affinity for actin filaments (Supplementary Fig. 4b). To query for F-actin barbed end binding, we utilized a seeded pyrene actin assembly assay to measure relative rates of elongation. Protein binding to assembling barbed ends usually modulates their elongation rates^11,31^, but we failed to detect any change in the presence of LCRs (Supplementary Fig. 4c). In support of these bulk biochemical assays, imaging of fluorescently labeled SNAP-LCR shows very minimal colocalization with actin filaments via TIRF microscopy (Fig. 4f; −27 and −23 min). Thus, LCR appears to have a very low affinity to the sides or the barbed ends of relaxed actin filaments.

Since Pxl1 and zyxin localize to actomyosin contractile structures, we hypothesized that these LCRs bind to an element common to both contractile actin networks. We reconstituted the core contractile machinery base on previously developed protocols^32,33^. An F-actin network was assembled from actin monomers, α-actinin, and SNAP-LCR, within buffer that contained 0.5% methylcellulose to crowd the network to a PEG-passivated glass coverslip surface. Individual actin filaments are mobile, but dynamics arrest as filaments elongate and are cross-linked by α-actinin into a network of mixed polarity bundles (Fig. 4b; 0 min). When the network reached an appropriate density, we flowed in a fresh mixture containing the initial concentrations of SNAP-LCR and α-actinin, the critical concentration of actin monomers (0.1 μM) to prevent network disassembly, and prepolymerized myosin II filaments (Fig. 4b; 30 min). Myosin II activity on F-actin generates local stresses and deformations that remodel and contract the network. Gently curved bundles become taut, with many breaking over time (Fig. 4b; 31 min and Supplementary Video 5). These failures usually occurred at bundle junctions (“T-junctions”) and more rarely along the length of a single filament or bundle (~10%). After rupture, bundle portions recoil and then compact into asters in which actin filaments are compressed, bent and severed^32^. Thus, the myosin-driven contraction of a reconstituted F-actin network occurs with a build-up of both tensile and compressive stresses on the actin filaments that drive filament buckling and breaking^32^.

Whereas LCR does not localize well to the networks initially, consistent with the ‘bulk’ F-actin sedimentation assays (Supplementary Fig. b), after the addition of myosin II LCR accumulates on a subset of the network structures most likely to be under high contractile tension (Fig. 4c-k; Supplementary Videos 6 and 7). The majority (~90%) of myosin II-induced breaks in the network occurred at T-junctions (Fig. 4c,d). Actin filaments at these junctions become highly distorted at sites of high tension and the actin filament(s) fail at or very near the junction point. In addition to being highly distorted, the change in the bundle thickness that occurs at T-junctions results in a discontinuity in bundle mechanics, which may result in the accumulation of stresses at the junction point. LCRs associate an average of 30 sec before T-junction failure and then immediately dissociated from the filament after breaking (Fig. 4c-e and Supplementary Video 5). Additionally, significantly more LCR binds to T-junctions that do not break compared to other random sites along actin bundles (Fig. 4d). To get a sense of the affinity of LIM domains for sites of stressed F-actin, we measured the residence time of LCR single molecule fluorescence on T-junctions. Since filament breaking results in LCR dissociation, this measurement represents a lower bound for the off rate, which must therefore be greater than 0.033 s^-1^. An assumption that the protein association rate is between 10^6^ – 10^7^ M^-1^ s^-1^ ^34^ suggests that LCR affinity for strained actin filaments is on the order of 1-10 nM. Significantly lower forces are needed to generate local strains by bending F-actin than by pulling on its long axis^35,36^, and we speculate that T-junctions that do not break may be enriched in highly curved actin filaments that are strained enough for LCR binding to occur.

Compared to T-junctions, myosin-induced breaks along the F-actin bundle are much rarer. However, when these breaks occur, SNAP-LCR(Pxl1) accumulation on the bundles appears dynamic with most localization occurring within 5 minutes prior to breaking at t=0 min (Fig. 4f,g). While the exact localization site varies between breaking events, SNAP-LCR signal is detected for an average of 60% of the 5 minutes prior to bundle break (Fig. 4h). LCR is also observed along control bundles that do not break throughout the course of the movie, but for only an average of 20% of the 5 minutes (Fig. 4h). LCR(Pxl1) only associates to the network after myosin addition, remaining on filament portions for several minutes prior to breaking at t=0 min (Fig. 4h).

LCR also co-localizes with myosin II, especially in areas with highly deformed F-actin networks (Fig.4i-k and Supplementary Video 7). While SNAP-LCR is detected above background on isolated myosin II (Supplementary Fig. 4d), LCR binding is ~5-fold higher when myosin is localized on the F-actin network, suggesting this colocalization is a result of LCR binding to myosin-induced actin filament deformations rather than directly to myosin (Fig. 4h). Line scans of a bundle over time show that the majority of LCR localization is independent of myosin localization (Supplementary Fig. 5). This further indicates that colocalization of LCR and myosin II along the network is due to LCR binding to actin filament deformations instead of myosin II directly (Fig. 4i,j). In actomyosin contraction assays that lack the cross-linker α-actinin, myosin drives contraction of the F-actin network into asters, which are dense clusters of actin and myosin II. Previous work has shown that these so-called asters are sites where filaments are buckled and broken due to compressive forces^32^. We observe that SNAP-LCR localizes to these asters, suggesting that LCR localization does not require the cross-linker α-actinin and localizes to highly compacted actomyosin (Fig. 4k). These data show that myosin II-generated forces on actin filaments is necessary and sufficient to drive LIM localization to actin filaments.

### Polymerization-generated stress is sufficient for LIM localization to actin

To determine whether LIM localization can occur by alternate means of applying force to actin filaments, we employed a well-established reconstitution assay that serves as a model for actin-based motion^37,38^. Here, 2 μm polystyrene beads are coated with the Arp2/3 complex nucleation promoting factor pWa (Fig. 5). Mixing the beads with actin monomers, Arp2/3 complex, and capping protein drives the assembly of a branched network of short actin filaments at the bead surface^39^, which can be visualized by imaging of fluorescently labeled actin. In the expanding actin shell, the outer network continuously stretches as it is displaced outward by continuous assembly of new actin at the bead surface. Forces generated by actin polymerization result in the buildup of circumferential tension along the outer part of the actin shell, resulting in network tearing that breaks the symmetrical shell. After symmetry breaking, the so-called comet tail drives directed bead motion^37^. Previous work has demonstrated that capping protein (CP) facilitates symmetry breaking, as short-capped filaments are more effective in force generation^40–42^.

**Figure 5.**
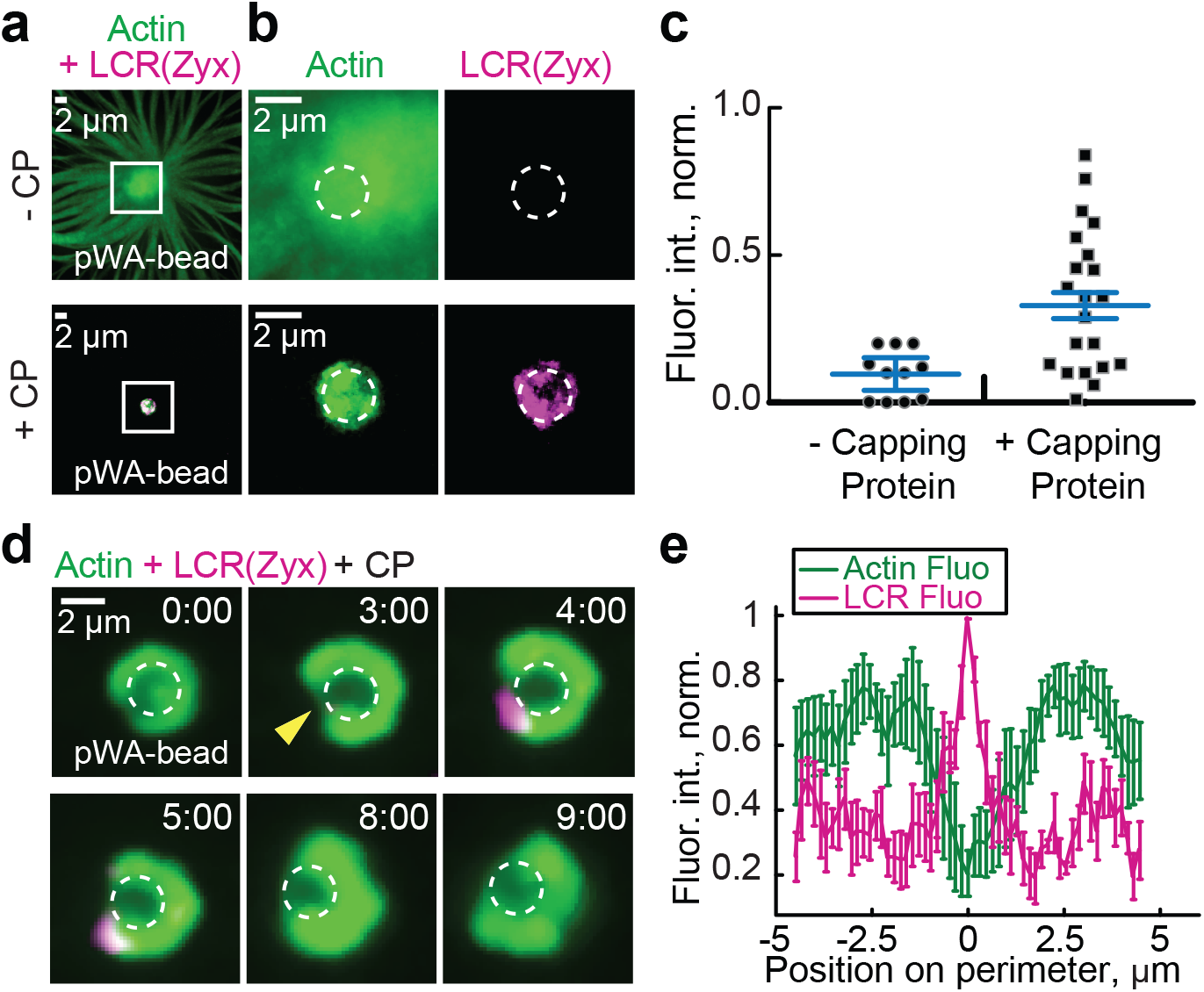
SNAP-LCR(zyx) localizes to branched F-actin networks during symmetry breaking. (a-c) LCR(zyx) localizes to capped F-actin networks in motile bead assays. (a) Confocal images taken from a motile bead assay, where the Arp2/3 complex activator pWa is electrostatically bound to the surface of polystyrene beads. The beads are mixed with 4 μM actin monomer (5% Alexa-488 labeled), 100 nM Arp2/3 complex, 12 μM profilin, and 400 nM LCR(zyx) in polymerization buffer in the presence or absence of 200 nM capping protein. (b) Zoomed images from the white boxed region of (a). White circle traces the beads. (c) LCR fluorescence on the actin networks in (a) in the absence or presence of capping protein. (d-e) HALO-LCR(zyx) localizes to symmetry breaking events in motile bead assays. The pWa beads are mixed with 2 μM actin monomer (5% Alexa-488 labeled), 100 nM Arp2/3 complex, 100-200 nM LCR(zyx), and 42 nM capping protein. (d) Confocal time-lapse of a representative symmetry breaking event. White circle indicates the beads, and the yellow arrow shows where symmetry breaks. (e) Linescans of LCR (magenta) and actin (green) fluorescence on the circumference of the F-actin network from reactions where beads break symmetry. Linescans were aligned by setting the peak LCR intensity to position zero.

In the absence of capping protein, large amounts of actin are assembled from the bead that elongates away from the bead surface. While the overall actin signal is much higher in the absence of capping protein, LCR binds only slightly above background detection (Fig. 5a-c). Conversely, in the presence of capping protein, we observed a wide distribution of LCR binding to the shell. A subset of beads has high LCR intensity associated with the actin shell (~50%), while others display levels of LCR binding similar to controls without capping protein (Fig. 5c). In confocal sections taken through the center of beads in the presence of capping protein, LCR localizes where the actin signal is weakest (Fig. 5d,e). We speculated that this is because symmetry begins to break at these sites and the F-actin network thins as it is strained. To look more closely at this phenomenon, we took confocal time-lapse movies of beads going through the process of symmetry breaking. We observed LCR localizing most intensely to the actin shell during the period of most rapid straining as the shell ruptures (Fig. 5d and Supplementary Video 8).

## DISCUSSION

Here we show that the mechanism by which zyxin is recruited to SFSS is through binding of its LCR exclusively to mechanically strained actin filaments. We identify 17 proteins from six different LIM domain protein classes with LCRs that localize to SFSS, indicating that this force-sensitive interaction may function as an input into diverse cellular processes. While SFSS are a particular feature within adherent fibroblasts, mechanical stresses are ubiquitous within the actin cytoskeleton. Force-sensitive biochemistry is inherent to mechanical regulation of the cytoskeleton and, we suspect, also a means for transmitting information about the mechanical status of the cell to the nucleus. The tension in SFs tends to reflect the mechanics of the environment in which cells are embedded. Cells growing in rigid matrices or within tissues that are being stretched build F-actin networks under increased tension^20,43,44^. Recent work has shown that matrix mechanics controls the nuclear localization of the LIM protein FHL2^45^. Interestingly, a majority of SFSS-binding LIM proteins display nuclear shuttling^46^, suggesting a model by which LCR binding to stressed F-actin networks blocks nuclear import. Our data raises the possibility that detection of cytoskeletal mechanics by LCRs from FHL, zyxin, tes, prickle and ajuba classes may underlie regulation of diverse transcriptional pathways they are part of, including YAP/TAZ, Hippo, p21 signaling and planer cell polarity. While much work has demonstrated these to be mechanically regulated^43,45,47,48^, our work implicates interactions between the LCR with strained actin filaments in diverse cytoskeletal assemblies as a potential mechanism. We anticipate that understanding the details of how and why diverse LIM domain containing proteins differentially localize to strained actin filaments at focal adhesions, cell-cell adhesions and the actin cytoskeleton will yield insight into the regulation and architecture of these mechanotransduction pathways (Fig. 6a).

**Figure 6.**
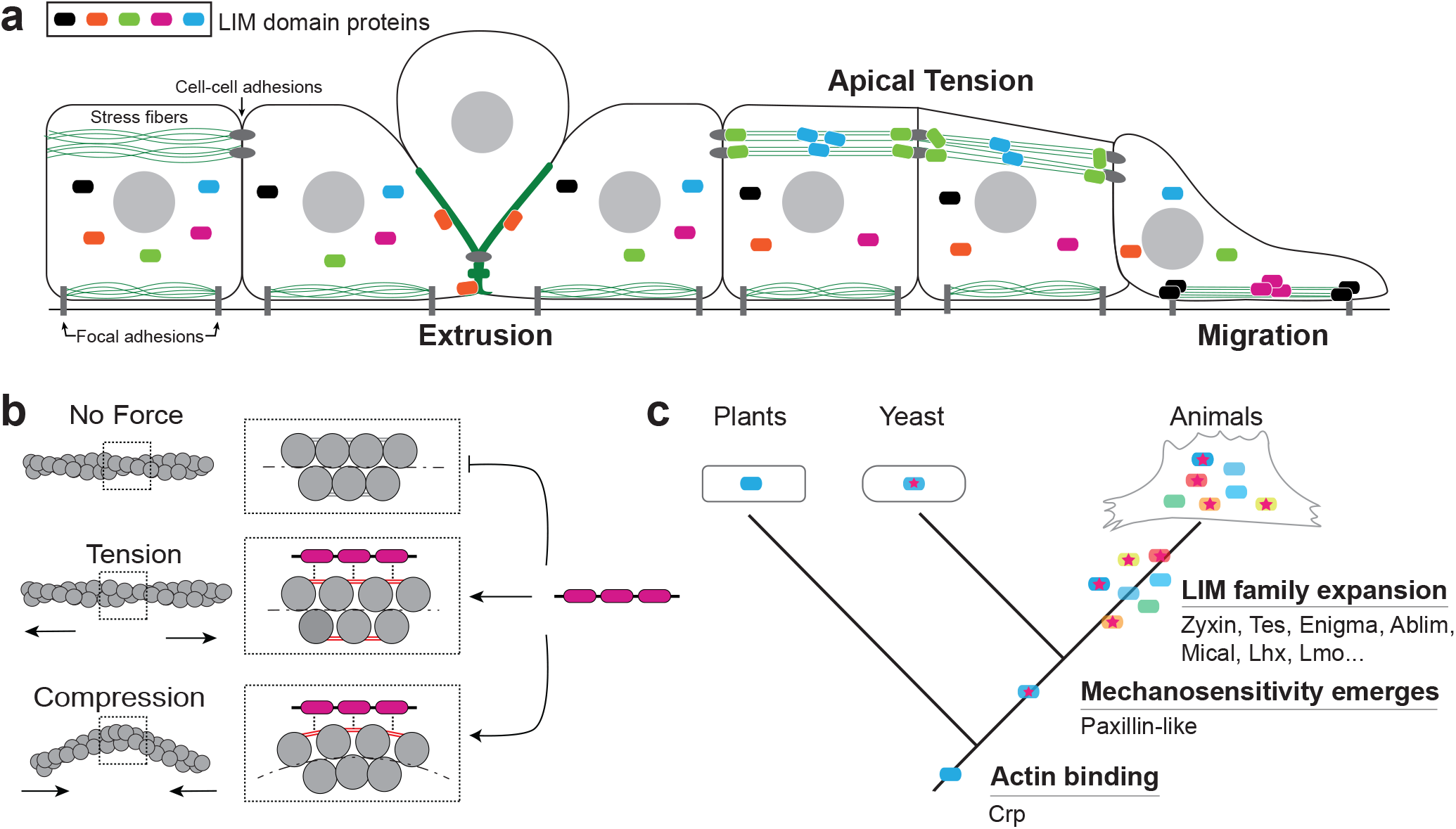
Model. (a) Cartoon of an epithelial tissue layer with different examples of mechanical stress that cells experience (extrusion, apical tension, and migration). LIM domains bind the different mechanically stressed F-actin networks. (b) Actin filaments experiencing different forces (tension or compression). The box indicates the zoomed in diagram showing the bonds between actin subunits. The grey lines are low stresses, while the red lines are high stresses. The LIM domain proteins likely localize to the regions of high stress. (c) Phylogenetic tree showing emergence of strained actin filament binding by paxillin class of LIM domain proteins and later expansion of LIM family in the metazoan stem lineage.

Our *in vitro* data indicate that the LCR can be recruited to highly tensed or compressed actin filaments, suggesting that these two distinct force-induced filament conformations may expose a similar actin filament structure that has high affinity for LCR (Fig. 6b). The maximum distortion that may occur within a highly bent filament before it breaks is estimated to be approximately 1.5A of displacement/subunit^35^. This relatively modest strain is enough to result in a conformation recognized by LIM domains. An attractive hypothesis is that stress exposes a site within the actin filament that is weakly recognized by each LIM domain. Our data show that tandem domains connected by linkers of a precise length, each contribute to binding a strained-induced feature on an actin filament. We suspect the linker length may act as a ruler that positions individual LIM domains to optimally bind a stress-induced feature on the actin filament (Fig. 6b).

We show that this force-sensitivity is found in fission yeast LIM protein Pxl1. The fact that Pxl1 both localizes to the contractile ring in fission yeast and to SFSS in animal cells suggests that myosin II-induced strained actin filament conformations are a common feature in contractile networks. Despite large evolutionary distances, this interaction has been conserved, indicating that there is significant selective pressure to maintain it. Actin is one of the most highly conserved proteins in Eukaryotes with 90.4% amino acid sequence identity between fission yeast and mammals, so it is not surprising that the LCR of Pxl1 from fission yeast binds to a strained F-actin structure in mammals. The oldest LIM domain protein found in plants and animals, CRP^25^, binds and bundles actin filaments via its LIM domains in the absence of mechanical stress^49–51^. We hypothesize that duplication and divergence of an ancestral CRP-like LIM domain resulted in a modification to its actin binding mechanism that favored a strained conformation of F-actin (Fig. 6c). The other core components of the contractile machinery, myosin II and α-actinin, have not been found in plants but are clearly present in the unikonts^52,53^ and may have appeared around similar times. We hypothesize that the emergence of contractile F-actin machinery coincided with, or required proteins that could report on the stresses present there. The oldest SFSS binding class from our screen appears to be paxillin^23^, which is involved in mechanical homeostasis of contractile networks in yeast^28^ and mammals^19^. F-actin strain sensing via LIM may have been co-opted by other signaling pathways later on during the LIM family expansion that originated in the stem lineage of metazoan (Fig 6c)^23^.

Zinc finger proteins have diverse functionality from regulation of cell cycle, transcription and protein folding through interactions with DNA, lipid and proteins^54^. Our data demonstrate that mechanically stressed actin filaments are an additional substrate for a subset of zinc finger proteins. The extent to which mechanical forces may regulate interactions with other known substrates is an opportunity to be explored. The use of the actin filament itself as a force sensor, or mechanophore, within the actin cytoskeleton is a particularly attractive one as means to control mechanotransduction pathways. Both its abundance and the different types of force (twist, compression, extension) that can be sensed could provide a wealth of control of mechanotransduction pathways. Moreover, our work suggests the possibility of other biomolecules that exclusively bind to mechanically stressed filaments that may not be have been isolated by traditional biochemical studies.

## Materials and Methods

### Cell culture and transfection

NIH 3T3 fibroblasts (American Type Culture Collection, Manassas, VA) and mouse embryo fibroblasts (MEFs) were cultured in DMEM media (Mediatech, Herndon, VA) and supplemented with 10% fetal bovine serum (HyClone; ThermoFisher Scientific, Hampton, NH), 2 mM L-glutamine (Invitrogen, Caarlsbad, CA) and penicillin–streptomycin (Invitrogen). Zyxin^(-/-)^ and zyxin^(-/-)^+EGFP-zyxin MEFs cells were a gift of Mary Beckerle’s laboratory (University of Utah, Salt Lake City, UT) and have been described previously^55,56^. All cells were transiently transfected via electroporation before experiments using a Neon transfection system (ThermoFisher Scientific). 1-2 μg of DNA was used per transfection. Following transfection, cells were plated in 8-well μ-slide chambers (Ibidi USA, inc.) and imaged 12-24 hours later.

### Live-cell imaging and SFSS induction

Cells were grown in 8-well μ-slide chambers with culture media supplemented with 10 mM HEPES and maintained at 37 °C. Cells were imaged on an inverted Nikon Ti-E microscope (Nikon, Melville, NY) with a Yokogawa CSU-X confocal scanhead (Yokogawa Electric, Tokyo, Japan) and laser merge module containing 491, 561 and 642 nm laser lines (Spectral Applied Research, Ontario, Canada). Images were collected on Zyla 4.2 sCMOS Camera (Andor, Belfast, UK). A 405 nm laser coupled to a Mosaic digital micromirror device (Andor) was used to locally damage SF. A small area targeting a region of a SF was drawn in MetaMorph and illuminated by the 405 nm laser for ~5 seconds. Images were collected using a 60x 1.49 NA ApoTIRF oil immersion objective (Nikon). All hardware was controlled using MetaMorph Automation and Image Analysis Software (Molecular Devices, Sunnyvale, CA). Cells were imaged in the 491 and 561 channel at various time intervals ranging from 2 – 20 seconds.

### Image processing and analysis for LCR Screening Assay

SFSS were analyzed by measuring the ratio of the mean fluorescence of transiently transfected mCherry-tagged LCR to the stably integrated EGFP-zyxin fluorescence as so: *Ratio = (LCR SFSS – LCR background) / (zxyin SFSS – zyxin background)*. Several images of the cell culture media region without cells was captured in each channel before an imaging experiment to obtain the background signal from camera noise, media fluorescence. These images were averaged and subtracted. On a day of imaging, care was taken to keep imaging settings constant. Each new day, at least two different transfections of LCR(zyx) were used as a control to adjust for day to day differences in imaging conditions. Kymographs were generated with imageJ reslice function, or if stress fibers moved significantly, a python script written in imageJ to build kymographs from user-drawn line segments was used. Background was considered to be the location on the stress fiber where the SFSS eventually developed. A linescan was drawn across the kymograph (through the time axis) to generate a profile (Supplementary Fig. 1f-o). The signal immediately before SFSS was subtracted from both channels.

The maximum fluorescence signal was found and averaged with the two adjacent points to obtain the enrichment value for each channel. If there was no obvious signal for the LCR being tested, then the time point at which GFP-zyxin signal was maximal was used in measuring the mCherry-LCR signal. The enrichment of mCherry was divided by the enrichment of GFP-zyxin to obtain the fluorescence ratio (Supplementary Fig. 1f-o). n was considered to be a single SFSS from four or more cells for each construct.

### Statistical analysis

All experiments were repeated a minimum of three times. Cells presented in figures are representative samples of the population behavior. Error bars represent the s.d., s.e.m or 95% confidence interval and were computed in graphpad prism (version 8.0d, GraphPad Software, Inc., La Jolla, CA) graphing software. Statistical significance was determined using independent two-sample Student’s t-test of the mean to compare groups of data. Statistical significance is indicated by asterisks: (*) represents a *p-value < 0.05*; (**) represents a *p-value < 0.01*.

### Plasmid constructs for SFSS screens and protein purification

LIM domains can be roughly defined by the following sequence motif: [C][X]_2_[C][X]_13-20_[H][X]_2_[C][X]_2_[C][X] _2_ [C][X]_13-20_[C][X]_2_[C], where the zinc coordinating residue [C] is usually cysteine but less frequently H, D, or E, and X is any amino acid. To identify LIM domain containing proteins, the LIM hidden Markov model (HMM PF00412.22) from pFam was used to scan mouse, fission and budding yeasts proteomes using hmmsearch (EMBL-EBI). Primers were designed to PCR amplify LIM domains from cDNAs generated from mouse embryo fibroblasts or fission yeast cells. 15 nucleotide overlaps were included in primers for infusion cloning (Clontech, Mountain View, CA). Where this failed, or for generating synthetic constructs as in (Fig. 3e), synthetic gBlock DNAs from IDT (Integrated DNA Technologies, Inc.; Coralville, IA) were designed, optimizing codons for expression and to reduce DNA repeat sequences. If possible, eight amino acids on either side of the first and last zinc coordinating residues were included in the LCR clones for the screen. Several initial LCR clones localized strongly to the nucleus. To ensure that LCRs were cytoplasmic, a nuclear export sequence l (NES) of zyxin was included on the amino terminal end of the LCR clones. A vector containing CMV promoter, zyxin’s NES, BsmBI and BamHI sites, GGSGGS linker and mCherry. LCRs were cloned into the BsmBI/BamHI cut vectors. BsmBI removes its own recognition sequence, so no additional residues were added in the NES-LCR ORF. For purification constructs, zyxin (residues 381-572) and *S. pombe* Paxillin-like protein (Pxl1; residues 256-438) PCR products were either inserted into pET21a-MBP-TEV at EcoRI/HindIII or into pET21a-MBP-TEV-SNAP at XmaI/NotI with a flexible linker (GGSGGS) in the forward primer following the SNAP sequence.

### LIM protein expression in fission yeast

For expression in fission yeast, full length sequences were cloned into genomic integration vectors pJK210-41xnmt-[LIM domain protein]-GFP::*ura4*+ that targeted a ura4 region in cells containing a ura4-294 loss of function point mutant, and restored the ability to grow in the absence of uracil when successful integration had occurred. Successful integration was secondarily screened by PCR and sequencing of the ura4 genomic region.

### Protein purification and labeling

Zyxin and Pxl1 LCR proteins were expressed in *Escherichia coli* strain BL21-Codon Plus (DE3)-RP (Agilent Technologies, Santa Clara, CA) with 0.5 mM isopropyl β-D-1-thiogalactopyranoside for 16 h at 16 °C. Cells were lysed with an Emulsi-Flex-C3 (Avestin, Ottawa, Canada) in extraction buffer [10 mM HEPES (pH 7.4), 150nM NaCl, 0.02% NaN3, 0.1 mM DTT] with EDTA-free Protease Inhibitor Cocktail (Roche, Basel, Switzerland) and were clarified. The addition of 50 μM ZnCl_2_ to the extraction buffers and subsequent buffers during later protein purifications resulted in an increased protein yield. The extract was incubated for 1 h at 4 °C with amylose resin (New England Biolabs, Ipswich, MA), loaded onto a column, washed with extraction buffer, and the protein was eluted with 30 mM maltose. LCR constructs were dialyzed against SNAP buffer [20 mM Hepes (pH 7.4), 200 mM KCl, 0.01% NaN_3_, 10% glycerol, and 1 mM DTT. SNAP-tagged proteins were filtered on a Superdex 200 10/300 GL or Superose 6 Increase 10/300 GL column (GE Healthcare, Little Chalfont, UK) and then labeled with SNAP-surface-549 (New England Biolabs, Ipswich, MA) overnight at 4°C following the manufacturers’ protocols. LIM proteins were flash-frozen in liquid nitrogen and stored at −80 °C. Actin was purified from chicken skeletal muscle acetone powder by a cycle of polymerization and depolymerization and gel filtration (Spudich and Watt, 1971). Gel-filtered actin was labeled with Alexa 488 carboxylic acid succinimidyl ester on lysine residues^57,58^. Human α-actinin IV was expressed in bacteria and purified as described^59^. Skeletal muscle myosin II was purified from chicken breast and labeled with Alexa 647 as described. Arp2/3 complex was purified from calf thymus by WASp(VCA) affinity chromatography^60^. WASP fragment construct GST-human WASp pWA was purified by Glutathione-Sepharose affinity chromatography ^29^.

### High-speed sedimentation

20 μM Mg-ATP actin monomers were spontaneously assembled in [10 mM imidazole (pH 7.0), 50 mM KCl, 6 mM MgCl_2_, 1.2 mM ethylene glycol tetraacetic acid (EGTA), 50 mM dithiothreitol (DTT), 0.2 mM ATP, 50 μM CaCl_2_] for 1 hr to produce F-actin. A range of F-actin (0-10 μM) was then incubated with 0.5 μM LCR construct for 20 min at 25°C. The mixture was spun at 100,000 g for 20 min, and the supernatant and pellet were separated by 12.5% SDS PAGE, stained with Coomassie Blue, destained, and analyzed via densiometry with ImageJ.

### Seeded pyrene

Pyrene assembly assays were conducted in 96-well plates in an Infinite M200 Pro (Tecan Systems, Inc., San Jose, CA) fluorescent plate reader to measure the fluorescence of pyrene-actin (excitation at 367 nm and emission at 407 nm. For seeded assembly, a 5 μM stock of unlabeled Mg-ATP actin monomers was preassembled, followed by the addition of 100x anti-foam, 10x KMEI [500 mM KCl, 10 mM MgCl_2_, 10 mM EGTA, 100 mM imidazole (pH 7.0)], and a range of LCR(zyx) (0-2 μM). The final concentration of F-actin unlabeled seeds was 0.5 μM for each well. Actin monomers (20% pyrene-labeled) were added to another row of wells, and mixing the monomer row with the preassembled actin filament row initiated the reactions.

### In vitro contractility assay

Time-lapse TIRFM movies were taken using a cellTIRF 4Line system (Olympus, Center Valley, PA) fitted to an Olympus IX-71 microscope with through-the-objective TIRF illumination and an iXon EMCCD camera (Andor Technology, Belfast, UK). The bundled actin networks consisted of F-actin (1.5 or 3 μM, 10% Alexa 488 labeled), α-actinin (100-200 nM), LCR (50-200 nM), and polymerization TIRF buffer [10 mM imidazole (pH 7.0), 50 mM KCl, 1 mM MgCl_2_, 1 mM ethylene glycol tetraacetic acid (EGTA), 50 mM dithiothreitol (DTT), 0.2 mM ATP, 50 μM CaCl_2_, 15 mM glucose, 20 μg/ml catalase, 100 μg/ml glucose oxidase, and 0.5% (400 cP) methylcellulose]. Based on the glass and imaging conditions, the specific protein concentrations varied within a set range to create contractile networks. The protein mixture was transferred to a flow chamber, and the network was allowed to preassemble for 30-45 minutes. Simultaneously, monomeric myosin II was polymerized separately in a low salt buffer [10 mM imidazole (pH 7.0), 1 mM MgCl_2_, 1 mM ethylene glycol tetraacetic acid (EGTA), 0.3 mM ATP. To induce network contraction, polymerized myosin (50-150 nM) was flowed into the chamber. The flow mixture also contained the critical concentration of actin (0.1 μM) and the same initial concentrations of α-actinin and LCR. Images were then acquired at either 10 or 15 s intervals at room temperature.

### Bead symmetry breaking assay

Polystyrene microspheres (Polysciences, Eppelheim, Germany) were coated with GST-pWA^60^. Motile beads were imaged after 15 minutes of polymerization on an inverted microscope (Ti-E; Nikon, Melville, NY) with a confocal scan head (CSU-X; Yokogawa Electric, Musashino, Tokyo, Japan), 491, 561, and 642 laser lines (Spectral Applied Research, Richmond Hill, Ontario, Canada) and an HQ2 CCD camera (Roper Scientific, Trenton, NJ). Z-stacks were acquired, reconstructed and analyzed using ImageJ. Fluorescence ratios were determined using the central, single plane of motile beads. Background-subtracted fluorescence values were measured for each fluorescent protein in the comet tail region and in the protrusion region.

## AUTHOR CONTRIBUTIONS

J.D.W. conceived the project and designed and performed experiments described in Figures 1, 2 and 3. C.A.A., J.D.W. and C.S. designed and performed experiments described in Figures 4 and 5. D.R.K. and M.L.G. guided progress of the project. All authors contributed to data analysis and wrote the paper.

## ACKNOWLEDGMENTS

We thank Mary Beckerle and her lab for the GFP-zyxin fibroblasts. We thank Greg Alushin, Mary Beckerle and Clare Waterman for helpful discussions and exchange of preliminary data over the last several years. Authors acknowledge funding from NIH R01 GM104032 (M.L.G.), NIH R01 GM079265 (D.R.K.), and ARO MURI W911NF1410403 (M.L.G. and D.R.K.), NIH F32 GM122372 (J.D.W)

