## Supplementary Material for "Evolutionarily diverse LIM domain-containing proteins bind stressed actin filaments through a conserved mechanism"

Supplementary Figures 1-5……………………………………………………...…………. 2-7

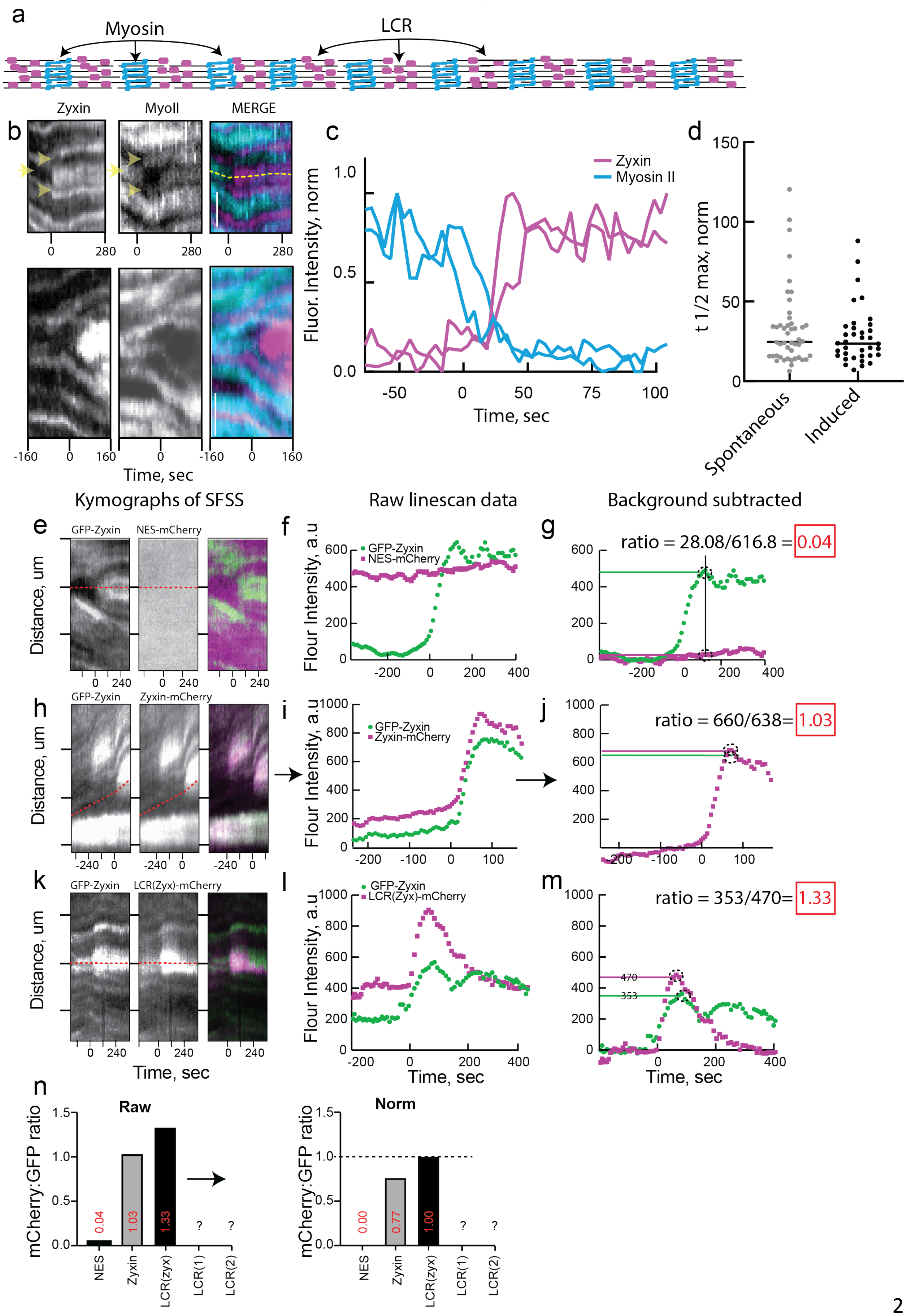

a
Myosin
LCR
b
Zyxin
MyoII
MERGE
Time, sec
c
Zyxin
Myosin II
Fluor. Intensity, norm
Time, sec
d
t 1/2 max, norm
Spontaneous
Induced
Kymographs of SFSS
Raw linescan data
Background subtracted
e
GFP-Zyxin
NES-mCherry
Distance, um
f
GFP-Zyxin
NES-mCherry
Flour Intensity, a.u
g
ratio = 28.08/616.8 = 0.04
h
GFP-Zyxin
Zyxin-mCherry
Distance, um
i
GFP-Zyxin
Zyxin-mCherry
Flour Intensity, a.u
j
ratio = 660/638= 1.03
k
GFP-Zyxin
LCR(Zyx)-mCherry
Distance, um
Time, sec
l
GFP-Zyxin
LCR(Zyx)-mCherry
Flour Intensity, a.u
Time, sec
m
ratio = 353/470= 1.33
470
353
Time, sec
n
Raw
mCherry:GFP ratio
0.04
1.03
1.33
?
?
NES
Zyxin
LCR(zyx)
LCR(1)
LCR(2)
Norm
mCherry:GFP ratio
0.00
0.77
1.00
?
?
NES
Zyxin
LCR(zyx)
LCR(1)
LCR(2)

**Supplementary Figure 1.** ***Diverse LIM domains from mammals localize to SFSS.***
(a) Cartoon Schematic of a stress fiber showing the periodic, complementary banding pattern of Myosin and LCR. (b,c) Myosin behavior in SFSS. (b) Kymographs taken from a time-lapse confocal movie of 3T3 cell with the heavy chain of myosin II fused on its amino terminus with mApple (Cyan) and stably integrated. Zyxin-mCherry transiently transfected. (Scale bar=2 μm). (c) Linescans through kymographs such as those shown in b, (yellow dotted line in merge) showing fluorescence intensity over time of myosin and zyxin. (d) Kinetics of Spontaneous and laser-induced SFSS. The time to reach half of the maximum fluorescence intensity ($t_{1/2}$) was calculated by taking linescans through kymographs as done in b and c. (e-m) General SFSS quantification workflow using mouse embryo fibroblasts with integrated GFP-zyxin. Kymographs and corresponding linescan data generated from SFSS with the mCherry-zyxin as a positive control (e-g), a negative control consisting of just a nuclear export signa tagged with mCherry (h-j), and the LCR being assayed, here the LCR from zyxin (LCR(zyx)) is shown (k-m). (f,I,l) Original uncorrected linescan data generated from the dotted red line on kymographs. (g,j,m) Background subtracted linescans. The signal before SFSS development is subtracted. From the background subtracted data, we determined the peak of each curve (dotted circles) and averaged the maximum with the two adjacent data points to determine the construct enrichment at the site. The enrichment of the mCherry construct was divided by the enrichment of GFP-zyxin to obtain the main metric, the mCherry:GFP fluorescence ratio. (n) For screening, the LCR(zyx) was assayed each different day so that constructs assayed on different days could be compared. The ratio of LCR(zyx)-mCherry:GFP-zyxin is always set to one.

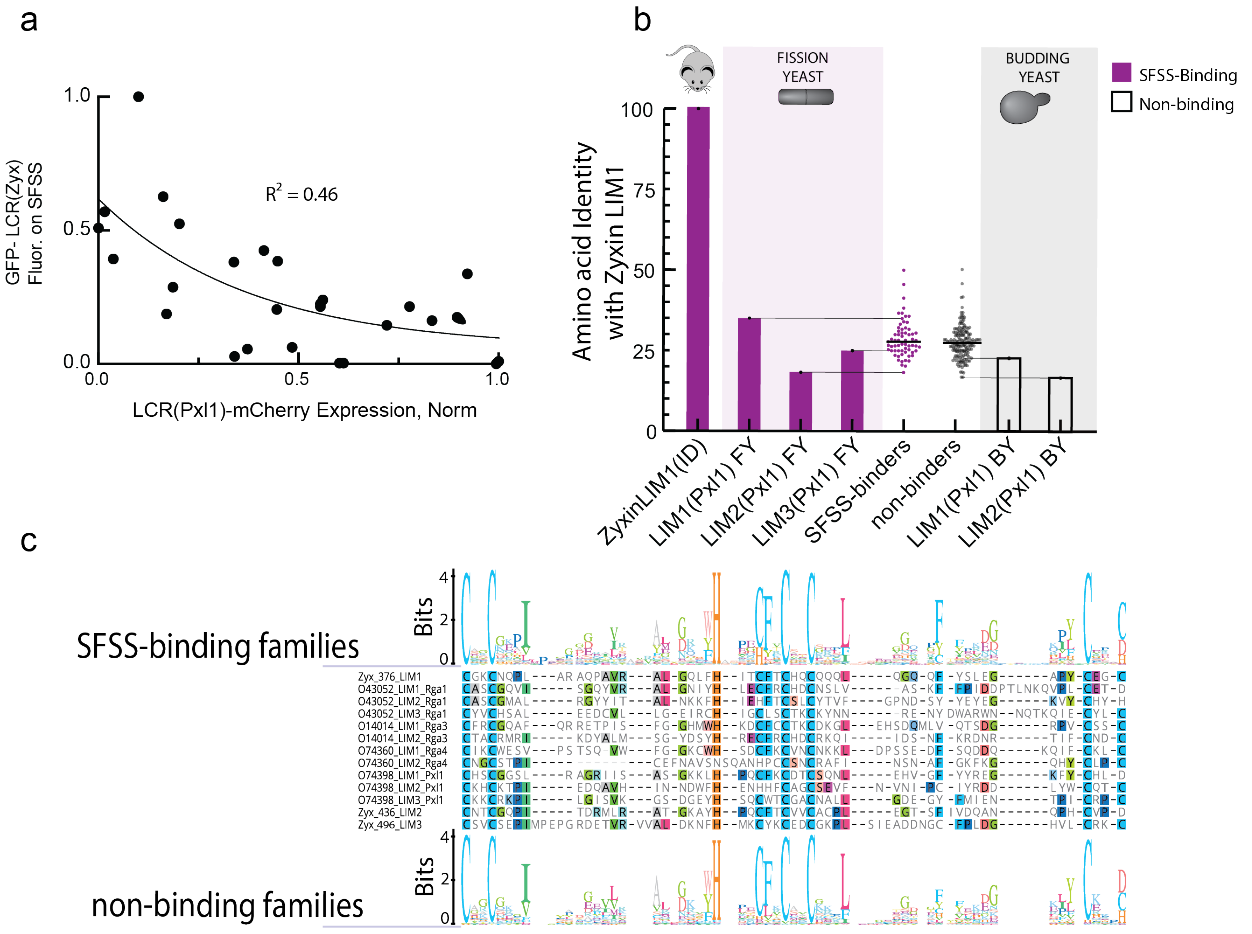


**Supplementary Figure 2.** *LCR from fission yeast Pxl1 and zyxin compete for SFSS and display similar kinetics.*

(a) The cytoplasmic signal of LCR(Pxl1) plotted against the LCR(Pxl1)-mCherry signal at the SFSS. Cytoplasmic signal was used as a proxy for expression level. (b,c) Diversity of LIM domains from fission yeast as compared with zyxin LIM domain 1, ID =Identity; zyxin LIM 1 is compared with itself here. (b) The three LIM domains from fission (FY) and budding yeast (BY) Pxl1 were aligned with all other mammalian and Yeast LIM domains and a distance matrix was generated. The distances between zyxin LIM 1 and all other LIM domains were plotted. (c) Sequence LOGOs of SFSS-binding and non-binding families.

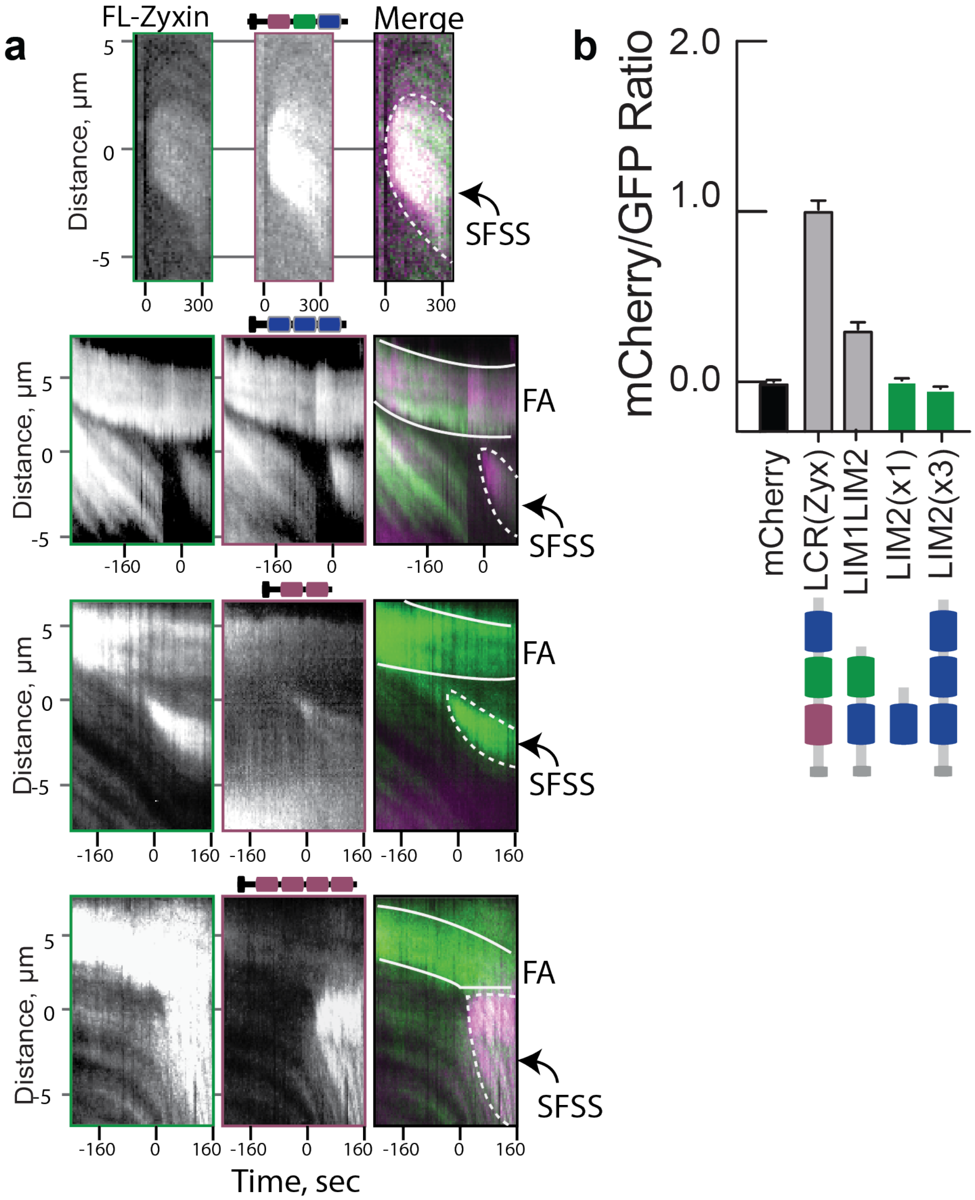


**Supplementary Figure 3.** *LCRs bind to SFSS through multiple, precisely spaced domains organized in tandem.*

(a) Representative kymographs taken from cells expressing various mutant constructs in MEFs with integrated GFP-zyxin. Top row is LCR(zyx)-mCherry, top-middle row is L3(x3), bottom middle row is L1(X2), and bottom row is L1(x4). FA=Focal adhesion. (b) SFSS binding by control LCR(zyx) compared to zyxin LIM2 constructs.

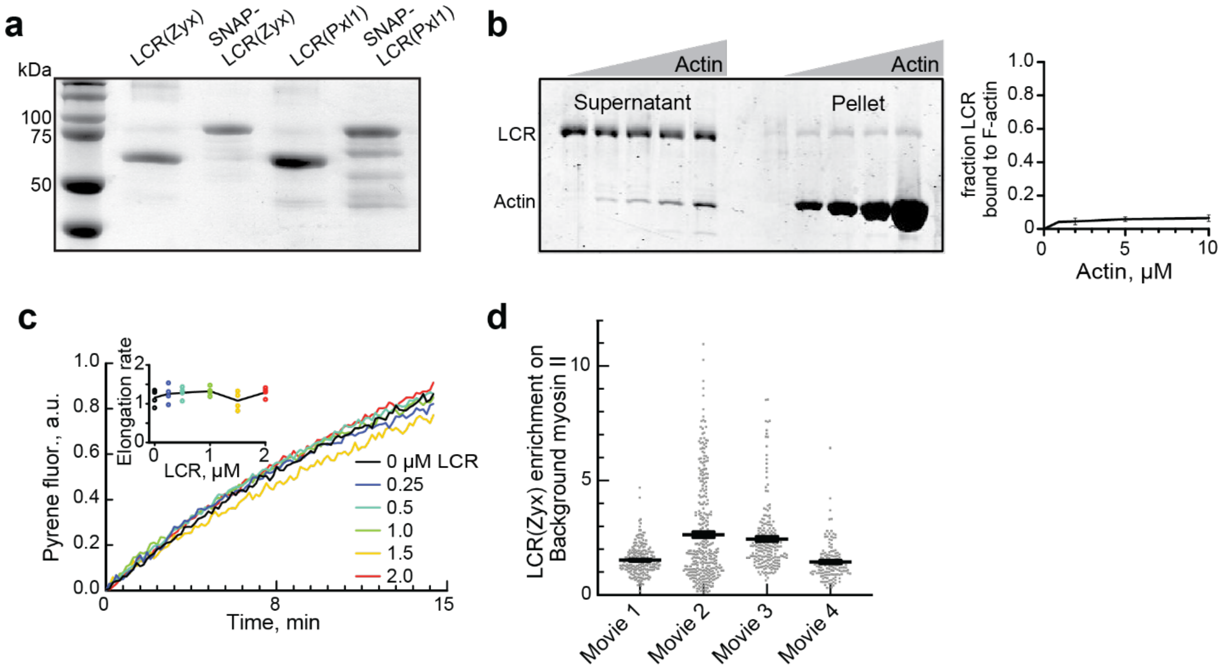


**Supplementary Figure 4.** *Purified LIM domains from fission yeast and mammals localize to stressed actin filament networks in vitro.* (a) SDS PAGE of Purified LCRs from mammalian zyxin and fission yeast paxillin Pxl1. (b) High-speed (100,000 x g) sedimentation assays of 0.5 µM LCR(zyx) with actin filaments preassembled from an increasing concentration (0, 1, 2, 5, 10 µM) of Mg-ATP actin monomer. (b, left) Coomassie stained gel of the supernatants and pellets. (b, right) Graph of the percent of total LCR(zyx) observed in the pellet vs the supernatant as a function of increasing actin concentration. (c) Seeded bulk actin assembly assay. Addition of 0.5 µM Mg-ATP actin monomers (20% pyrene labeled) onto 0.5 µM preassembled filaments in the presence of a range of LCR(zyx) concentrations (0-2 µM). (c Inset) Plot of initial slopes of the curves, used as a proxy for elongation rate. Assembly from spontaneous nucleation is not observed under these conditions, as there is no observable increase in pyrene signal in the absence of seeds (data not shown). (d) SNAP-LCR(zyx) fluorescence within myosin II puncta that were adhered to a PEG-passivated surface, but not associated with the F-actin network *in vitro* contraction assay described in Fig. 4. The average LCR fluorescence within the puncta was divided by the average local background signal to get an enrichment of LCR within Myosin puncta compared to background.

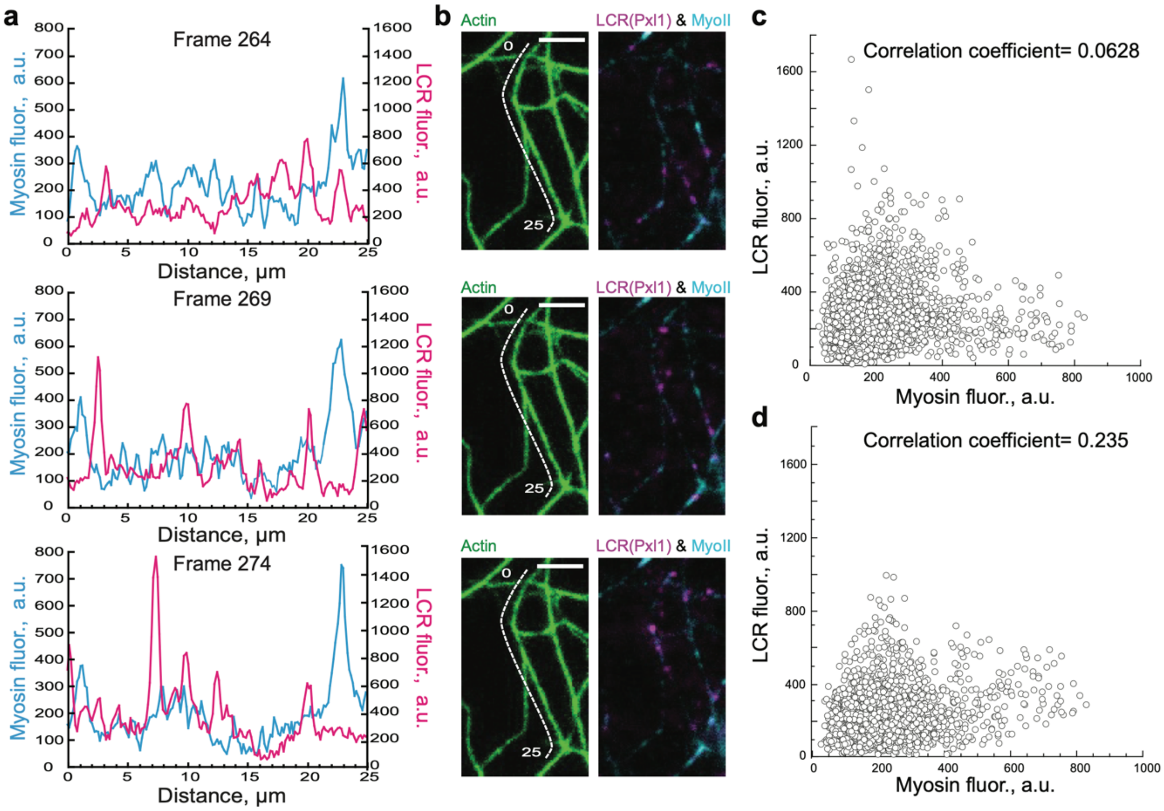


**Supplementary Figure 5.** *LCR(Pxl1) localization along reconstituted F-actin networks does not correlate with myosin localization.* (a-c) Quantitative analysis of LCR(Pxl1) and myosin localization along an F-actin bundle in the *in vitro* contraction assay described in Fig. 4. A preassembled network was formed with 3 µM actin, 200 nM α-actinin, and 20 nM SNAP-LCR(zyx), followed by the addition of 300 nM myosin to induce network contraction. The network was imaged every 15s. (a) Myosin (cyan, left y-axis) and LCR(Pxl1) (magenta, right y-axis) linescans taken along a single F-actin bundle over time. Linescans for frames 75s apart are shown here. (b) The corresponding images for the linescans in (a). White lines indicate the bundle starting at 0 µm and ending at 25 µm. Scale bar=5 µm. (c-d) Plots of LCR fluorescence vs myosin fluorescence for the bundle in (b) over 20 frames. (c) The Pearson's correlation coefficient was calculated for the relationship between LCR and myosin, 0.0628. (d) As a control, the LCR channel was rotated 90° to the right, and new values for the same ROIs in (c) were plotted against the same myosin values in (c). The Pearson's correlation coefficient is 0.235 for this plot.
